# Filamented Light (FLight) Biofabrication of Mini-Tendon Models Show Tunable Matrix Confinement and Nuclear Morphology

**DOI:** 10.1101/2025.02.03.636123

**Authors:** Hao Liu, Lynn Scherpe, Linnea B. Hummer, Jess Gerrit Snedeker, Marcy Zenobi-Wong

## Abstract

One hallmark of healthy tendon tissue is the high confinement of tenocytes between tightly packed, highly aligned collagen fibers. During tendinopathy, this organization becomes dysregulated, leading to cells with round-shaped morphology and collagen fibers which exhibit crimping and misalignment. The elongated nuclei in healthy tendons are linked to matrix homeostasis through distinct mechanotransduction pathways, and it is believed that the loss of nuclear confinement could upregulate genes associated with abnormal matrix remodeling. Replicating the cell and nuclear morphology of healthy and diseased states of tendon, however, remains a significant challenge for engineered *in vitro* tendon models. Here we report on a high throughput biofabrication of mini-tendons that mimick the tendon core compartment based on the Filamented Light (FLight) approach. Each mini-tendon, with a length of 4 mm, was composed of parallel hydrogel microfilaments (2-5 µm diameter) and microchannels (2-10 µm diameter) that confined the cells. We generated four distinct matrices with varying stiffness (7-40 kPa) and microchannel dimensions. After 14 days of culture, 29% of tenocytes in the softest matrix with the largest microchannel diameter were aligned, exhibiting an average nuclear aspect ratio (nAR) of 2.1. In contrast, 84% of tenocytes in the stiffest matrix with the smallest microchannel diameter were highly aligned, with a mean nAR of 3.4. When tenocytes were cultured on the FLight hydrogels (2D) as opposed to within the hydrogels (3D), the mean nAR was less than 1.9, indicating that nuclear morphology is significantly more confined in 3D environments. By tuning the stiffness and microarchitecture of the FLight matrix, we demonstrated that mechanical confinement can be modulated to exert control over the extent of nuclear confinement. This high-throughput, tunable platform offers a promising approach for studying the mechanobiology of healthy and diseased tendons and for eventual testing of drug compounds against tendinopathy.

## Introduction

Tendons are multicellular fibrous tissues that transmit force from muscles to bones, facilitating body movement.[1] Tendon tissue exhibits a well-organized hierarchical structure, primarily composed of intrinsic (core) and extrinsic (peritenon) compartments.[2,3] The intrinsic compartment consists of a well-organized fibrous extracellular matrix (ECM) and resident tendon cells, while the extrinsic compartment includes the immune, vascular, and nervous systems. Changes in the organization of the fibrous ECM and cellular morphology within the core intrinsic compartment is often closely associated with the onset of tendon disease.[4,5] Furthermore, these alteration in the intrinsic compartment can trigger a cascade of immune responses, with “cross-talk” between the intrinsic and extrinsic compartments being essential for mediating tendon regeneration and ECM remodeling homeostasis.[6] However, to date, we have yet to engineer tendon tissues that closely mimic the intrinsic compartment, limiting our understanding of tendon physiology and pathology.

Collagen type I is the primary extracellular matrix (ECM) component in intrinsic compartment of tendons, exhibiting highly aligned organization in its healthy state, which results in uniaxial tensile moduli of up to 1’000 MPa.[7] In healthy tendons, collagen fibers have been observed to have diameters ranging from 1 to 100 μm.[8,9] The single collagen fiber structures have lengths ranging up to centimeters, with an aspect ratio often exceeding 100:1. The spacing between these collagen fibers varies from a few hundred nanometers to several micrometers in a mechanical unloaded state.[10] However, in degenerative tendon diseases collectively referred to as tendinopathy, the organization of collagen fibers is significantly altered. The fibers are less tightly packed with loss of their parallel alignment in the core of tendinopathic tendons.[4,11] Additionally, there is a shift toward a more heterogeneous ECM composition, for instance, a significant increase in collagen type III. All of these changes in collagen organization and composition ultimately result in a significant decrease in mechanical properties of pathological tendons compared to healthy tendons.[12]

At the tissue and cellular level, tenocytes from tendinopathic tissue show increased proliferation and higher metabolic activity, with cross-talk to the extrinsic compartment associated with neovascularization, and pain-associated nerve ingrowth compared to healthy tendon.[2,13] The shift between healthy and tendinopathic cells is also associated with a change in nuclear morphology, from an elongated morphology with nuclear aspect ratio (nAR) often above 4.0, to a more rounded one (nAR close to 2.0).

The mechanical confinement imposed by the three-dimensional (3D) matrix and the associated nuclear morphology is linked to the ECM organization. Microarchitecture and material stiffness both influence the amount of mechanical confinement a cell will experience.[14] As the nucleus deforms, specific mechanotransduction pathways are activated, thereby influencing cell transcriptional behavior.[15–17] Therefore, developing tissue models that mimic the differences in extracellular matrix organization and nuclear morphology observed in intrinsic tendon compartment is crucial for advancing our understanding of tendon physiology and pathology.[12,18,19]

Current biofabrication techniques for engineered human tendon models provide cells with 3D scaffolds and topological cues to induce the anisotropic tissue organization seen *in vivo*. Methods such as electrospinning and 2D surface patterning have been shown to effectively promote cell alignment, tendon differentiation, and the upregulation of key tenogenic markers such as collagen type I and scleraxis.[20–22] However, these approaches have primarily focused on providing topological cues for cell alignment, with less emphasis on replicating the nuclear morphology within 3D environments that is imposed by mechanical confinement. There is a pressing need for engineered *in vitro* tendon models that can precisely mimic both the matrix organization and nuclear morphology of intrinsic tendon compartment.

Filamented Light (FLight) is a light-based biofabrication approach that enables the rapid and biocompatible encapsulation of cells within 3D anisotropic hydrogels.[23] FLight creates parallel, highly aligned microfilaments and interconnected high aspect ratio void spaces (microchannels) within the printed hydrogel. We previously showed that these ultrahigh-aspect-ratio (>500:1) microfilaments and microchannels, with diameters ranging from 2 to 20 μm, formed a highly porous hydrogel (∼50% porosity). The dimensions of these microchannels are close to the reported spatial size limits (∼3 μm) for cell migration, allowing for cell alignment and migration while simultaneously inducing nuclear deformation.[24]

In this study, we demonstrate the biofabrication of engineered *in vitro* tendon models (mini-tendons) that mimic the native intrinsic compartments using FLight technology. The hydrogel microfilaments closely resemble the dimensions and aspect ratio of native collagen fibers and effectively guide cell alignment in 3D. Additionally, we developed various FLight matrices with different stiffnesses and microchannel dimensions; these exert distinct mechanical confinements, ultimately resulting in different nuclear morphologies.

## Results

### 2.1 Matrix and Nuclear Morphology in Healthy and Unhealthy Tendons

To guide the interpretation of subsequent studies, cell and nuclear morphology of freshly harvested healthy and pathologic tendons were evaluated based on quantification of histological sections. Healthy and unhealthy tendon tissues exhibited differences in nuclear and ECM morphology (**Figure 1**, also **Figure S1**), consistent with previous reports.[13] Specifically, more elongated nuclei were found in healthy tendons as determined by nAR, defined as the long-to-short axis ratio of nuclei (**Figure 1b** and **Figure S1b**). The mean nAR in healthy tendons was found to be 6.2, compared to 2.6 in unhealthy tendons. In addition to nuclear morphology, a significant difference in cellularity was observed between healthy and unhealthy tendons. Unhealthy tendons exhibited an approximately 1.8-fold higher cellularity compared to healthy tissues (**Figure 1c**), which may be associated with the hypercellularity/hypervascularization that is typically observed in pathological tendons.[2] Furthermore, there was a notable discrepancy in the alignment of collagen type I fibers between the two groups, with the “aligned” defined as an angle between –10 and 10° from the orientation distribution peak. In unhealthy tendons, only 32% of collagen fibers were aligned, compared to 53% in the healthy tendons (**Figure 1d, Figure S1d**). This difference in collagen fiber alignment was also reflected in the nuclear alignment, where aligned nuclei are defined as those with their long axis oriented within a range of –10 to 10° from the peak of orientation distribution (**Figure S1c**). Approximately 45% of nuclei in healthy tendons were oriented along the long-axis direction of collagen fibers, compared to only 18% in unhealthy conditions (**Figure S1**). Overall, several pivotal metrics were identified through a comparative analysis of healthy and unhealthy tendons, including nAR, cellularity, and alignment of collagen fibers. Subsequent assessments of the effects of different FLight matrices on tenocytes were conducted with a focus on these critical factors.

**Figure 1.**
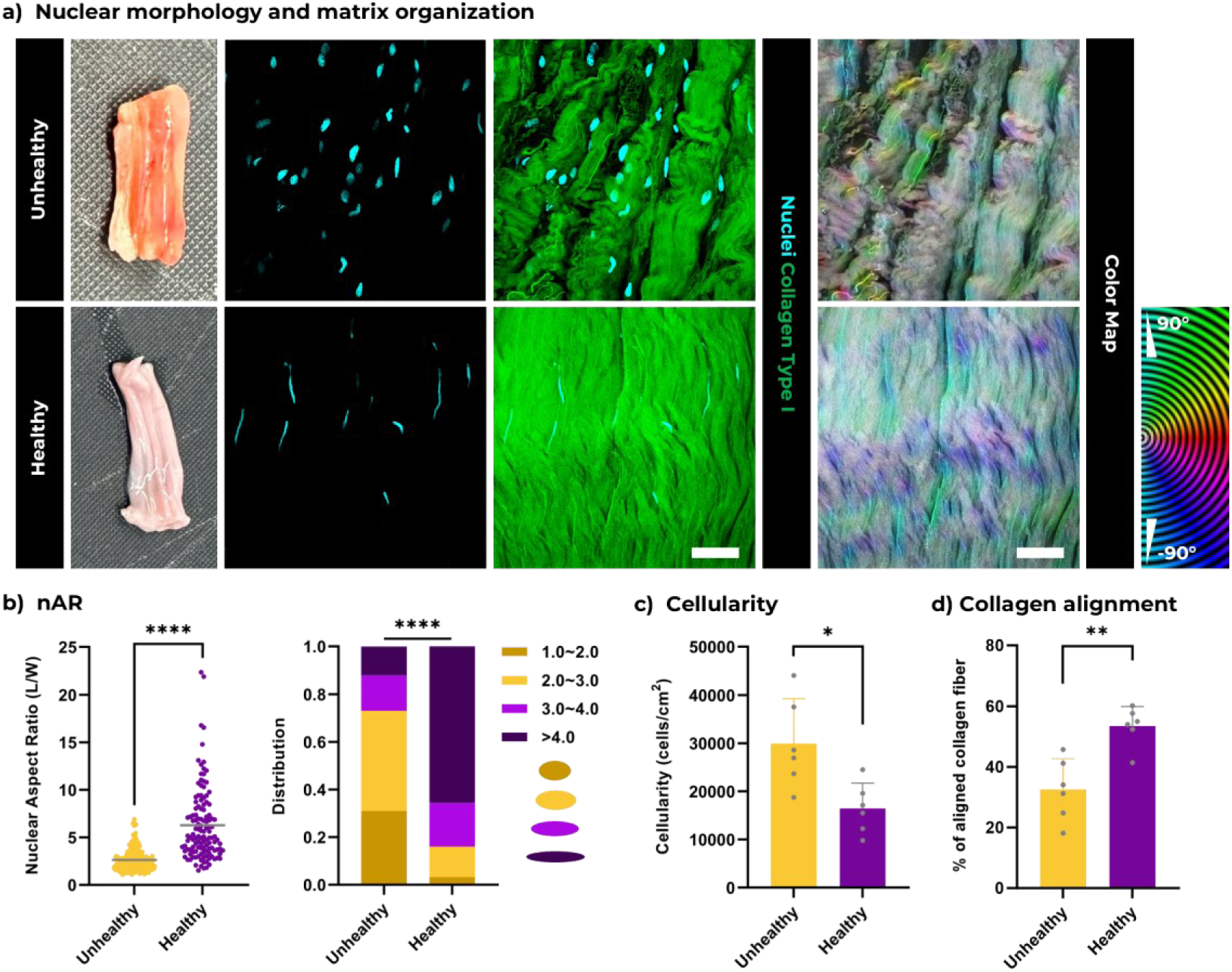
Nuclei morphologies in healthy and unhealthy human tendons. **a**) Photographs of healthy and unhealthy human tendons alongside representative confocal microscopy images showing cell nuclei and collagen type I. The right column depicts the orientation color map obtained from the OrientationJ plugin, corresponding to the confocal microscopy images. Scale bars: 20 µm. **b**) Nuclei aspect ratio (nAR) of cells and its distribution of nAR in healthy and unhealthy tendons (6 patients, dataset size>125). **c**) Cellularity comparison between healthy and unhealthy tendons. Additional details about the patients are available in the supporting information. **d**) Percentage of aligned collagen fibers in healthy and unhealthy human tendons. Collagen fibers are classified as aligned, characterized by an angle deviation from the peak distributed direction between –10 to 10°.

### 2.2 FLight Biofabrication of Hydrogel Matrices with Varying Microarchitectures and Mechanical Properties

FLight biofabrication of mini-tendon models is outlined in **Figure 2**. Tenocytes from a range of donors were expanded in 2D culture and then resuspended in a Gel-MA photoresin to create patient specific constructs (**Figure S2a**). A circular array was projected to create 96 cylindrical, mini-tendon constructs after exposure to 12 seconds of light (**Figure S2c, d**). The designed circles have a diameter of 1 mm and are equal to 37 pixels on projection images. The dimensions of each printed hydrogel construct were approximately 4 mm in length, measured along the direction of microfilaments, and 1 mm in diameter.

**Figure 2.**
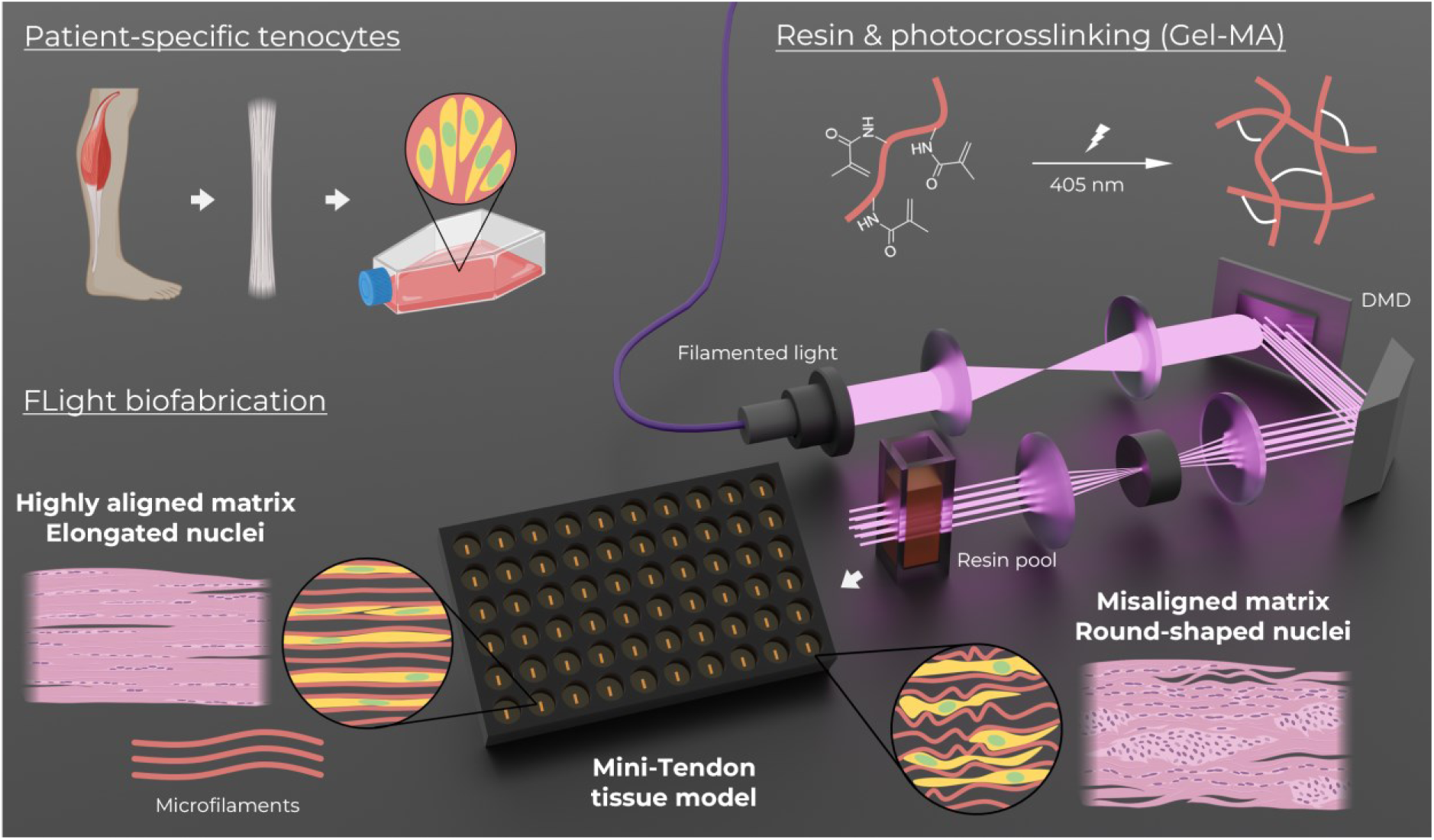
Schematic illustration of study design: This figure depicts the working principle of FLight (Filamented Light) biofabrication of the tissue-engineered tendon constructs (mini-tendon). Freshly isolated human tenocytes are expanded in 2D culture and mixed with Gel-MA photoresin to prepare the bioresin. Due to the optical modulation instability (OMI), the speckle pattern, characterized by local maxima and minima intensity distributions, is present in the projected laser beams, leading to filamentation of light.[51] Through modulation by a digital micromirror device (DMD), pre-designed geometric images are projected onto a resin pool. When the filamented light illuminates to an optically non-linear medium (i.e., (bio)resin), a self-focusing effect occurs, guiding the filamented light to photo-crosslink the (bio)resin into microfilaments. After the removal of uncrosslinked (bio)resin, microchannels are formed within the hydrogel constructs. These hydrogels, featuring various microfilaments and microchannels, create different matrix microenvironments (i.e., mechanical confinements) for the encapsulated tenocytes, leading to distinct cell/nuclei morphologies and diverse cellular behaviors. Up to 96 *in vitro* hydrogel constructs in millimeter scale are created within a matter of seconds. These constructs are then transferred to culture plates for subsequent incubation and analysis. Graphics were created in part using BioRender.

**Figure 3a** shows the microfilaments and microchannels of the four types of filamented matrices (M1-4) that were prepared using fluorescently labeled Gel-MA at varying polymer concentrations and light doses (**Table 1**). Notably, all hydrogel microfilaments were observed to be nearly perfectly aligned in the projection direction immediately following printing. However, the microfilaments in M1 (lowest light dose) exhibited a more disorganized morphology after washing out the uncrosslinked Gel-MA photoresin. M1 microfilaments were less aligned compared to M4 filaments (highest light dose, 10% polymer concentration), with only about 59% of microfilaments aligned within a range of –10 to 10° along the projection direction. In contrast, M2-4 exhibited an excellent alignment, with aligned microfilament ratios reaching 98% (Figure 3b). The main reason attributed to this microfilament disorganization is a lower degree of photocrosslinking of Gel-MA. A softer microfilament is less likely to maintain its alignment and physically confine a cell. In addition, as discussed in our previous work, reduced cross-linking between microfilaments at lower light projection doses may also contribute to the differences in confinement.[23] As evidenced by larger microchannel diameters, less crosslinking between microfilaments leads to separation of microfilaments and formation of a more disorganized morphology.

**Figure 3.**
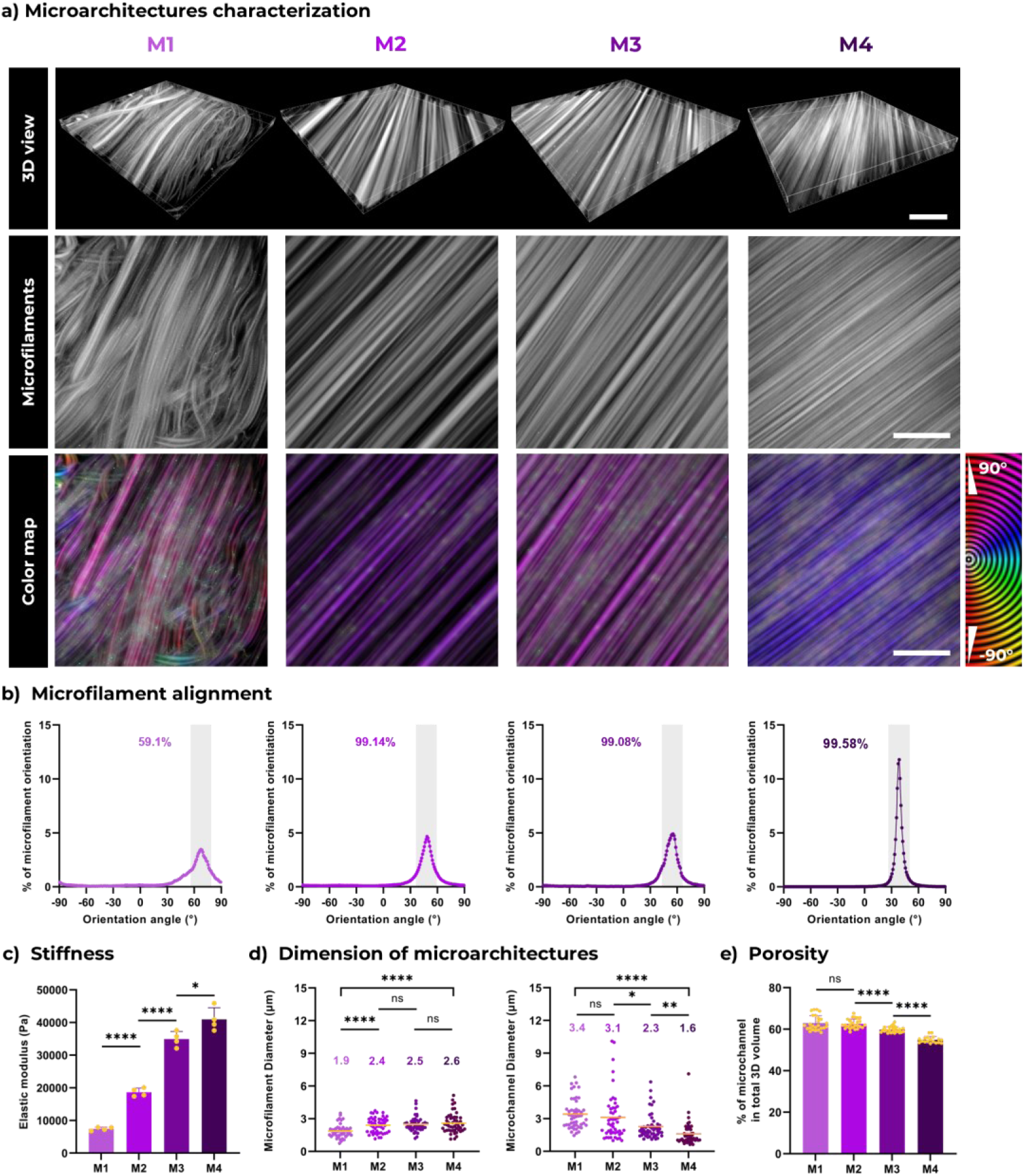
Characterization of microarchitectures in varying FLight hydrogel matrices. **a**) Representative confocal microscopy images showcasing microfilaments and microchannels across different hydrogel matrices (Matrix 1-4). The top row presents a 3D view of microarchitectures, generated from 3D reconstruction of Z-stack scans using Imaris 9.9.0 software. The middle row displays maximum-intensity projection images from Z-stack scans. The bottom row features an orientation color map obtained from the OrientationJ plugin, corresponding to the projection images. Scale bars: 50 µm. **b**) Distribution of microfilament orientation in various hydrogel matrices (Matrix 1-4). Gray areas highlight microfilaments classified as aligned, characterized by an angle deviation from the projection direction between –10 to 10°. Numeric labels represent the percentage of aligned microfilaments. **c**) Mechanical properties of hydrogel matrices evaluated using compression modulus (n=4). The compressions were applied in parallel to the microfilament direction. **d**) Analysis of microfilament and microchannel diameters across different FLight matrices (n=3, dataset size=50). The above values indicate the mean diameters. **e**) The ratio of microchannels in the total 3D hydrogel volume, indicating the porosity of the FLight matrices (n=3, dataset size>18).

**Table 1.** Compositions of photoresin and light dose for different FLight matrices.

| Entries | Polymer (Gel-MA)<br>Concentration<br>(w/v, %) | Photoinitiator<br>(LAP, w/v, %) | Light Dose<br>(mJ/cm<sup>2</sup>) | Compressive<br>Elastic Modulus<br>(kPa) |
| --- | --- | --- | --- | --- |
| M1 | 5 | 0.05 | 350 | 7.4 |
| M2 | 5 | 0.05 | 600 | 18.6 |
| M3 | 5 | 0.05 | 850 | 35.1 |
| M4 | 10 | 0.05 | 400 | 41.2 |

The stiffness of the filamented hydrogels increased in accordance with light-dose and Gel-MA concentration, with mean compressive moduli of 7.4, 18.6, 35.1, and 41.2 kPa for M1-4, respectively (Figure 3c). Measurements of the microarchitecture dimensions within M1-4 revealed that thicker microfilaments and narrower microchannels were found in M4 (Figure 3d). Compared to M1, where the mean microfilament diameter was 1.9 µm, the mean diameter in M4 increased to 2.6 µm. Conversely, the diameter of the microchannels exhibited a notable decrease, from 3.4 µm in M1 to 1.6 µm in M4. Furthermore, the ratio of 3D volume occupied by microchannels, also often defined as the porosity of the 3D hydrogel matrix, decreased to ≈ 55% in M4 compared to ≈ 63% in M1 (see Figure 3e).

### 2.3 Cellular and Nuclear Morphologies in Different Filamented Hydrogel Matrices

Healthy tenocytes from patients were encapsulated at a consistent density (0.3 million/mL) to examine their behaviors and morphologies in various filamented hydrogel matrices (M1-4). High cell viability (> 92%) was observed in all construct formulations and at all time points post-FLight biofabrication, with no significant difference in viability across all matrices after 0, 3, 7, and 14 days of culture (**Figure S3a, b**). Nuclear morphology (nAR) and cytoskeletal alignment (filaments actin, F-actin) were tracked at different time points (Figure 4). As previously reported, the encapsulated tenocytes gradually migrated into the microchannels from the original encapsulation site during the first few days of culture, aligning accordingly.[23] A loss of alignment of microfilaments was observed over time in M1, while in M2-4, microfilaments remained highly aligned even after 2 weeks of culture (Figure 4a, **Video S1**). The nAR of encapsulated tenocytes increased in all matrices at day 3 compared to the initial post-encapsulation phase (day 0). However, the nAR showed a progressive increase with 14 days of culture in M2-4, but not M1 (see Figure 4b). Given the major differences in matrix organization and nuclear morphology between M1 and M4, comparisons were focused on these two matrices (M1 vs. M4). After 2 weeks of culture, the distribution of nAR showed that nuclei in M4 were more elongated compared to those in M1, which were more round-shaped (**Video S2**). Specifically, 49.9% of the nuclei in M4 had an nAR > 3.0, while only 10.4% of the nuclei in M1 did (for nAR > 4.0: 28.3% in M4 vs. 3.2% in M1, Figure 4c). Of note, M1 showed a 5-fold higher cellularity than M4 on day 14, despite the same cell-seeding density. Similar trends were confirmed in the cell proliferation studies, where a significantly higher ratio of Ki-67 positive tenocytes was observed at days 3 and day 7 in M1 (**Figure S3c, d**), indicating more proliferation. Additionally, the cytoskeletons of cells in M1 were less aligned: only 19.1% of F-actin was aligned along the long axis of hydrogel microfilaments, whereas 83.6% of the F-actin was highly aligned in M4 (Figure 4e). The above results demonstrated that different FLight matrices lead to variations in nuclear morphology and proliferation.

**Figure 4.**
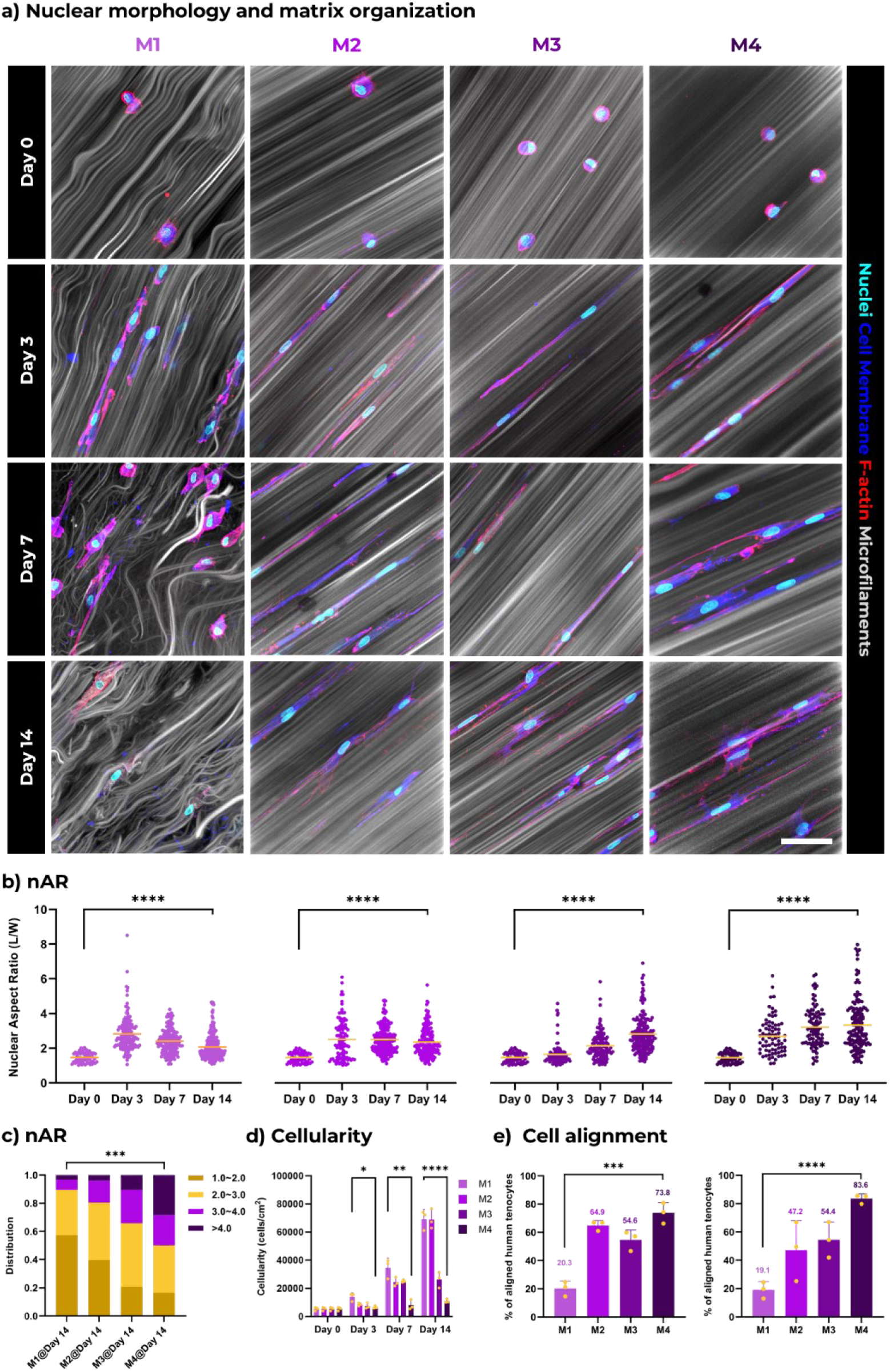
Nuclei morphologies in mini-tendon constructs. **a**) Representative confocal microscopy images showing human tenocytes in different mini-tendon constructs after 0, 3, 7, and 14 days of culture. The column labels indicate the specific matrices (as depicted in Figure 3) in which the cells were encapsulated. Images are maximum-intensity projections, scale bar: 50 µm. **b**) Analysis of nuclei aspect ratio (nAR) in human tenocytes at different culture durations, and **c**) the distribution of nAR in Matrix 1-4 following 14 days of culture (n=3, dataset size >22). **d**) Cellularity comparison within Matrix 1-4 at day 0, 3, 7, and 14. **e**) Proportion of aligned human tenocytes in different mini-tendon constructs, measured after 7 days (left) and 14 days (right) of incubation. The aligned cells are characterized by an angle deviation of f-actin from the projection direction between –10 to 10° using F-actin staining.

More importantly, after encapsulating tenocytes from other donors, we observed a misaligned matrix organization and lower nAR in M1 compared to those nuclei found in M4 (**Figure S4a-c**). In M1, tenocytes from all donors exhibited a significant increase in cellularity after 14 days of culture compared to those encapsulated in M4 (**Figure S4d**). Moreover, all different tenocytes in M4 were observed to be highly aligned along hydrogel microfilaments but not those in M1 (**Figure S4e**). All above results suggested that these variations are independent of individual patient differences and reliant on the properties of the FLight hydrogel matrices.

Another aspect not to be overlooked is the significant differences in the morphology of microfilaments between M1 and M4 over time. The progressive loss of alignment of microfilaments in M1 may have partially resulted from remodeling of the Gel-MA hydrogel,[25,26] as evidenced by the gene expression profile of matrix metalloproteinases (MMPs) in encapsulated tenocytes (**Figure S5**). The differences in transcript levels of the MMP3, MMP9, and MMP13 genes in M4 exhibited expression patterns similar to those observed in healthy tendons.[27,28] In contrast, a pattern similar to that found in pathological tendons was observed in M1. A significant increase in MMP9 and MMP13 expressions was observed in M1 compared to M4. These genes are known to be involved in the degradation of ECM, particularly gelatin and collagen, and are associated with excessive matrix turnover in pathological conditions.[19] Conversely, a marked decrease in MMP3 expression was observed in M1, perhaps reflecting the association between MMP3 gene expression and matrix remodeling in the context of tendon regeneration.[5,19] Clinical studies have also shown that the MMP3 expression is downregulated in tendinopathy compared to control groups.[29]

### 2.4 Different Mechanical Confinements in 2D and 3D environments

To understand the importance 3D confinement in a filamented hydrogel has on nuclear morphology, tenocytes were post-seeded onto the surfaces of M1 and M4 (2D) and compared to cells encapsulated within the FLight hydrogels (3D). The hydrogel microfilaments on the hydrogel surface merely provided topological cues to guide cell alignment while exerting minimal confinement. A mathematical model based on previous research was first constructed to predict the mechanical confinements exerted on tenocytes as a function of the biophysical parameter provided by 3D hydrogel matrices.[30] The predicted mechanical confinements applied to encapsulated tenocytes (3D) and 2D cultured tenocytes depended on matrix stiffness and microchannel diameter **(Figure S6)**. Predicted mechanical confinements also increased with matrix stiffness but were inversely correlated with microchannel diameter. In the 3D environment, the M4 is predicted to impose approximately 1.6-fold higher mechanical confinement on encapsulated cells compared to M1. When comparing 2D and 3D environments, cells within the 3D matrix are predicted to experience 3.4 times higher confinement than cells seeded on the 2D FLight hydrogel.

Next, we evaluated the predictions regarding differences in mechanical confinement by immunofluorescence staining of nuclear lamina and nuclear morphology. The distribution and assembly of lamin A/C, a nuclear lamina protein that is important in nuclear stability and force sensing, has been employed as an indicator of mechanical force transferred from matrix to cell.[31,32] As shown in **Figure S7**, in the 3D environment, cells in the M4 exhibited a higher nAR and increased lamin A/C expression. Comparing 2D and 3D environments, tenocytes on the surface of the M4(M4_2D) did not show similar nuclear morphology and lamin A/C expression to those presented in M4. These results suggest not only a difference in mechanical constraints between the M1 and M4 matrices but also, consistent with the predictions of the mathematical modeling, a significant difference in confinement between 2D and 3D environments.

Finally, we investigated whether differences in nuclear morphology resulting from mechanical confinement lead to the activation of specific mechanotransduction pathways. Among the various mechanosensitive pathways, Yes-associated protein 1 (YAP) was selected, given its potential role in responding to 3D confinement and mediation of several critical cellular behaviors such as proliferation and matrix remodeling.[15,33,34] The spatial distribution of YAP within tenocytes was analyzed under four conditions (i.e., M1_2D, M4_2D, M1 and M4) using confocal microscopic scanning and 3D image reconstruction (Figure 5a and **Figure S8**). The YAP spatial distribution and translocation were assessed by the volume of YAP positive staining divided by the nuclear volume. Tenocytes located on the hydrogel surface or presented within hydrogel matrices (i.e., 2D vs. 3D environments) exhibited significant differences in YAP nuclear localization. The highest nuclei-YAP overlap was observed in the tenocytes encapsulated in M4 – the condition exposed to the highest mechanical confinement (Figure 5c**-e**). In addition, we examined whether this relationship between nuclear morphology and YAP distributions was present in our human tissue samples. A significantly higher overlap ratio of YAP and nuclei has been found in healthy tendons compared to unhealthy tendons, indicating more nuclear translocation of YAP within these elongated nuclei (**Figure S9**).

**Figure 5.**
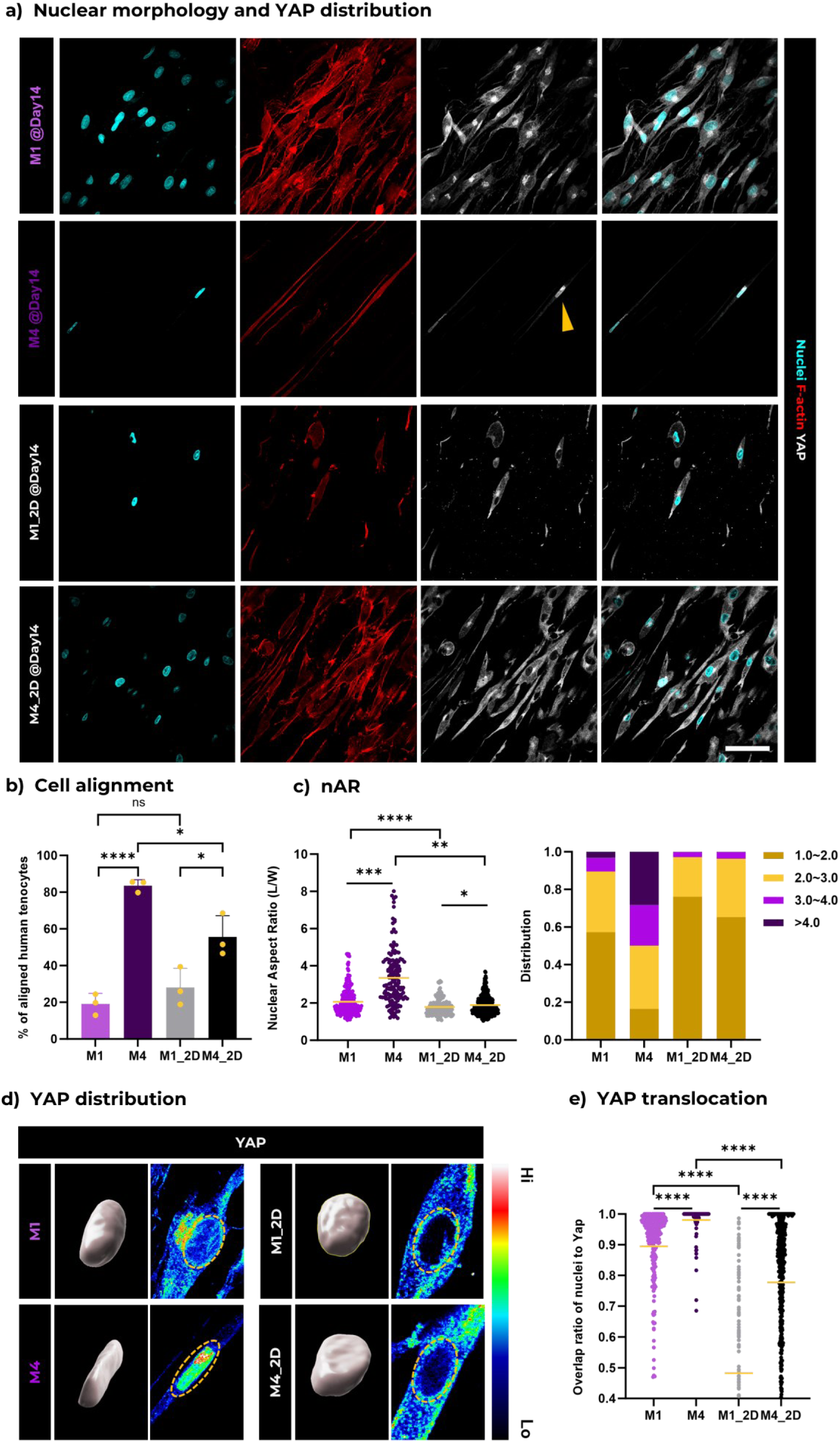
Nuclei morphologies and YAP nuclear activation of tenocytes under different mechanical confinements. **a**) Representative confocal microscopy images of human tenocytes within various mini-tendon constructs (Matrix 1 and 4), and on the 2D surface of FLight hydrogel constructs (Matrix 1 and 4) after 14 days of culture. Scale bar: 50 µm. **b**) Quantification of aligned f-actin in human tenocytes. F-actin alignment is defined by an angle deviation from the peak distributed direction between –10 to 10°. **c**) Analysis of the nuclei aspect ratio (nAR) in or on different hydrogel matrices, along with the distribution of nAR following 14 days of culture (n=3, dataset size >109). **d**) Representative images showing the nuclear morphologies of tenocytes and the YAP intensity distribution map. The yellow dashed line indicates the nuclei location. The 3D reconstruction of the nuclei was performed using Imaris 9.9.0 software, with relative intensity displayed on a rainbow scale. **e**) Measurement of nuclear volume and **f**) the overlap ratio of nuclei to YAP. A ratio of 0 indicates an absence of YAP signal within the nuclei, whereas a ratio of 1 denotes the presence of YAP signals encompassing the entire nuclear volume (n=3, dataset size >109).

In conclusion, we have identified that FLight matrices can be engineered to elicit biologically relevant differences in confinement and nuclear morphology to affect important mechanotransduction pathways.

## Conclusion & Discussion

In this study, we demonstrated the rapid biofabrication of 3D filamented hydrogel constructs as engineered tendon models using patient-specific tenocytes. A range of mini-tendons with varying mechanical properties and microarchitectural dimensions were created. We observed the best cell alignment and highest nAR in M4 – the stiffest matrix with the smallest microchannels. In contrast, softer matrices with larger microchannels exhibited less aligned tenocytes with more rounded nuclei. These differences in nuclear morphology were at least partly attributed to the different degrees of mechanical confinement imposed by the filamented matrices. Based on the mathematical model and the expression of nuclear lamin A/C, the confinement in the 3D matrix was significantly greater than in the 2D environment. Mechanical confinement in 3D environments was determined by a combination of matrix stiffness and microchannel diameter, resulting in the highest confinement within the M4 matrix. Finally, the differential nuclear localization of YAP suggests that these variations in nuclear morphology can lead to the activation of specific mechanotransduction pathways. The use of FLight biofabrication to create engineered *in vitro* tendon models makes it possible to closely replicate the matrix organization and nuclear morphologies observed in the native tendon core compartment.

However, the tuning of mechanical confinement in the current FLight process is contingent upon the light doses and polymer concentrations, which simultaneously affect the hydrogel stiffness, dimension of architectures, and even porosity. Future research could explore how individual matrix parameters, such as microchannel diameter, specifically influence nuclear morphology and activation of mechanotransduction pathways. Given that FLight technology for producing filamented hydrogel constructs involves light modulation instability and optical self-focusing effects in the non-linear medium, there is the opportunity to finely tune individual matrix properties by adjusting optical settings and photoresin properties. For example, spatial light modulators could be used to modulate speckle noise (e.g., coded phase mask),[35] or the refractive index could be matched to adjust self-focusing effects, as with the use of iodixanol.[36,37]

Given the flexibility of the FLight biofabrication and its suitability for a range of photocrosslinkable biomaterials, we can expect the development of more biomimetic tendon models in the near future. Recent advances in the photocrosslinking of natural biomaterials have demonstrated the use of ruthenium (Ru) and sodium persulfate (SPS) as photoinitiators to crosslink collagen or volumetrically print decellularized extracellular matrix (dECM) under 450 nm wavelength light, achieved through tyrosine crosslinking.[38,39] This opens new avenues for creating filamented 3D hydrogel constructs using natural matrix materials or their combinations, which could enhance the biomimetic properties of filamented matrices by incorporating chemical compositions closely resembling native tendon tissues. By tuning the chemical composition of the matrices (e.g., the ratio of collagen type I to type III in photoresins) and their corresponding matrix organization, there is potential to advance tissue matrix models towards mimicking both healthy and diseased tendons.[2,4] Most importantly, the multi-material/multi-cellular capability of FLight biofabrication may pave the way for remodeling the intrinsic-extrinsic compartments observed in tendons.[6,40,41] For example, after biofabricating the intrinsic tendon compartment, a secondary FLight projection can be applied to adjacent regions using a bioresin specifically tailored for the extrinsic compartment, thereby creating a sophisticated multicellular tendon model. By precisely tuning the properties of the intrinsic compartment, researchers can systematically explore how the extrinsic compartment respond, ultimately enabling a deeper understanding of the dynamic interactions and cross-talk between the intrinsic and extrinsic compartments.

The development of dynamic hydrogel networks using synthetic biomaterials could enable enhanced spatiotemporal control over tendon matrices. Specifically, enzyme-degradable/crosslinkable and photo-responsive hydrogels offer dynamic matrix properties for spatiotemporal tuning of mechanical confinement.[42–44] For example, modifications to hyaluronic acid with both methacrylate and *o*-nitrobenzyl (*o*NB) acrylate have been achieved. UV irradiation in the absence of a photoinitiator results in the degradation of *o*NB and the softening of the hydrogel, whereas in the presence of a photoinitiator, the photocrosslinking of methacrylate groups induces the stiffening of the hydrogel.[45] When combined with FLight biofabrication, these biomaterials show great potential for creating advanced matrix platforms for the study of nuclear morphology and cellular behavior in dynamic microenvironments.

Finally, while more elongated nuclei were observed in the highly confined microenvironment, the nAR still differed from those found in healthy native tendons (nAR > 6). Achieving a higher nAR may require further optimization of matrix properties, but the effects of mechanical loading or stimulation must also be considered. Previous studies have shown that cyclic stretching of *in vitro* tendon constructs and *ex vivo* explants induces cell alignment and upregulates gene expression associated with tendon repair, maturation, and ECM deposition.[46,47] In addition, the shear forces between hydrogel microfilaments and encapsulated cells under cyclic stretching is expected to activate specific mechanotransduction pathways such as Piezo1, enhancing tendon mechanical properties through increased collagen crosslinking.[48]

## Experimental Section

### Synthesis of Gel-MA and fluorescent-labeled Gel-MA

Gel-MA was synthesized following a previously reported protocol.[23] The degree of substitution (DS) was determined using ^1^H NMR (Bruker Ultrashield 400 MHz, performing 1024 scans) in D2O (Apollo Scientific). The integration signal for Gel-MA lysine (2.90–2.95 ppm) was compared against the signal for unmodified gelatin lysine (2.90–2.95 ppm). Phenylalanine signal (7.2–7.5 ppm) was used as an internal reference.

To prepare fluorescent-labeled Gel-MA, fluorescein-5-isothiocyanate (FITC) or rhodamine B isothiocyanate (RBITC) were conjugated to the Gel-MA. Briefly, 10% w/v of Gel-MA was dissolved in 100 × 10^−3^ M sodium bicarbonate solution. Subsequently, a 0.1% w/v of FITC-dimethylformamide or rhodamine solution was added, and the mixture was stirred at 40 °C for 6 h in the dark. Following the reaction, the mixture underwent dialysis against deionized water for 4 days at 30 °C, eliminating unreacted monomers, and was finally freeze-dried to yield the fluorescent-labeled Gel-MA.

### Cell culture

Human tendon samples and tenocytes were kindly provided by Prof. Jess G. Snedeker. These samples and cells were collected from patients undergoing knee or shoulder transplant surgeries according to a protocol approved by an ethics committee [BASEC 2020-01119; Tendon disease: Elucidating the molecular mechanisms that control tendon homeostasis and pathogenesis]. Healthy tendons and tenocytes were isolated from the semitendinosus or gracilis tendons, while unhealthy tendons were obtained from the biceps. Detailed patient information related to these samples is available in the supporting information (**Table S1**).

Tenocytes were expanded and cultured in T75 flasks with low glucose Dulbecco’s Modified Eagle Medium (DMEM; Gibco, 31885) + 10% v/v fetal bovine serum (FBS; Gibco, 10270) +1% v/v penicillin-streptomycin (Gibco, 15140) supplemented with 50 µg mL^-1^ L-ascorbic acid (TCI, A2521). Tenocytes were passaged at 80-90% confluency and detached using 0.25% Trypsin/ethylenediaminetetraacetic acid. To culture cell-laden hydrogel constructs, the samples were incubated in standard humidified conditions with 5% CO2 at 37 °C.

### Photoresin and Bioresin Preparation

Photoresin was prepared by dissolving the freeze-dried Gel-MA in 1× phosphate-buffered saline (PBS) solution at 50 °C for 30 min, subsequently mixed with photoinitiator, lithium phenyl-2,4,6-trimethylbenzoylphosphinate (LAP) to achieve a final concentration of 0.05% w/v LAP. The photoresin was filtered by 0.2 µm filter (SARSTEDT AG, Filtropur S 0.2), to sterilize it and remove potentially scattering particles. The human tenocytes were resuspended in photoresins at a concentration of 0.3 million cells mL^-1^ to prepare the bioresin.

### FLight biofabrication

(Bio)photoresin was prepared as mentioned previously and transferred into sterilized cuvettes (Thorlabs, CV10Q14F). Before being loaded into the custom-designed FLight3D printer with a wavelength of 405 nm, the cuvettes were kept at 4 °C for 15 minutes, facilitating the thermal gelation of Gel-MA. Projection images were generated using Affinity Photo (Serif Europe Ltd., Affinity Suite 1.9), set to a consistent resolution of 1024 × 768 pixels. These images were in 8-bit grayscale format, with pixel values reaching 255 to signify 100% light intensity (≈65 mW cm^-2^). The width of each pixel in the projection image was equal to 27 µm. Subsequently, the images were saved as PNG/TIFF files and uploaded to the FLight3D printer’s integrated software. Projection durations were determined based on the required light doses for varying experiments (**see Table S2**). Following FLight biofabrication, the uncrosslinked (bio)photoresin was washed away using 1×PBS prewarmed to 37 °C. The projected hydrogel constructs were carefully removed from the cuvette using a sterile spatula and then placed into 6-well plates filled with 1×PBS (for acellular constructs) or culture medium (for cell-laden constructs).

### Mechanical testing

For the compressive tests, cylindrical models were prepared, each with a diameter of 5 mm and a height of 4 mm. The hydrogel samples were tested by unconfined uniaxial compression using Texture Analyzer (Stable Micro systems, TA.XTplus). The apparatus included a 500 g load cell and a flat plate probe with a diameter of 15 mm. To ensure complete contact between the hydrogel samples and the plates, a preload of 0.2 g was applied. Samples were compressed to a final strain of 30%, processing at a rate of 0.01 mm s^-1^. The elastic compressive modulus was calculated by linear fitting of the initial linear region (0.5-5%) of the stress-strain curve. These tests were repeated four times at 25 °C to ensure consistency.

### Mathematical modeling

The mathematical model was established based on previously reported work.[30] The model posits that cells, initially encapsulated with spherical morphology in filamented hydrogel matrices, actively attempt to adjust their morphology to migrate and elongate within microchannels. The model simplifies the mechanical confinement as forces applied by unilateral hydrogel microfilaments on deformed tenocytes (as a semi-elliptical soft body), which are functions of cell stiffness,[49] ECM stiffness, cell dimension, and channel diameter. All parameters used in this model are listed in **Table S2**.

### (Immuno)fluorescence staining and confocal microscopy imaging

Cell-laden hydrogel constructs were washed with pre-warmed 1× PBS three times after 0, 3, 7, or 14 days of culture and then fixed in 4% paraformaldehyde (PFA) for 20 min at 25 °C. For human tendon tissues, fixation was done using 4% PFA at 4 °C for 6 hours, followed by embedding in paraffin for sectioning in the sagittal plane. Subsequently, the human tendon tissues and hydrogel samples were treated with 0.2% v/v Triton-X100 in PBS for 30 min for permeabilization, prior to being blocked with 1% w/v bovine serum albumin (BSA) in PBS for 1 h at 25 °C. These samples were then incubated with primary antibodies diluted in 1% w/v BSA-PBS for 12 h at 4 °C. Next, the samples were washed three times with 1× PBS, incubated with 1:500 diluted secondary antibodies, 1:1000 diluted Hoechst 33342, and pre-prepared phalloidin-tetramethylrhodamine B isothiocyanate working solution (0.13 µg mL^−1^, P1951) in 1% w/v BSA-PBS for 2 h at 4 °C. Imaging was executed with a 20×/60× objective lens on a confocal laser scanning microscope (CLSM; Olympus, Fluoview FV3000). Z-stack scanning was acquired from the 100 µm depth of constructs at 0.5-3 µm step sizes. The antibody details are provided in the supporting information (**Table S3**).

### Confocal microscopy image analysis and 3D reconstructions

For nuclear aspect ratio and cellularity measurements, the resulting Z-stack scans were transformed into Maximum-intensity projection images utilizing the ‘Z-project’ function in Fiji software. To segment nuclei, DAPI images underwent initial processing with a threshold setting. Subsequently, the ‘analyze particles’ function within Fiji was applied to determine the aspect ratio and cell count, incorporating a filter to include only sizes of interest larger than 50 pixels (n=3). Cellularity was evaluated by calculating the ratio of the number of nuclei to the area of the image (n=3).

To assess the alignment of microfilaments and F-actin, fluorescent-labeled Gel-MA was used to fabricate FLight hydrogel constructs. The CLSM imaging protocol was selected to match the excitation/emission wavelengths of the fluorescent molecules. The acquired image stacks were initially processed into projections and segmented as previously outlined. Subsequently, the ‘OrientationJ’ plugin (available at http://bigwww.epfl.ch/demo/orientationj)[50] was employed to evaluate the orientation and distribution of the aligned microfilaments. Similar methods were applied for analyzing alignment of F-actin, where confocal microscopy images from the mini-tendon samples were processed in the same manner (n=3).

To characterize the dimensions of microarchitectures, the acquired fluorescence images were segmented, and a line perpendicular to the microfilaments was drawn using the ‘Straight’ function. The diameters of both microfilaments and microchannels were then calculated from the data presented in the ‘Plot Profile’ panel in Fiji.

For the analysis of nuclear volume and YAP co-localization ratio, Z-stack scans were first reconstructed in 3D using Imaris software (Oxford Instruments, ver. 9.9.0). The ‘Surface’ function was employed to perform separate 3D reconstructions for the nuclei and YAP fluorescence channels. By default, the surface resolution was set at 0.418 µm, with a threshold value of 650, and no filters were applied. Using the ‘Vantage’ function, 2D plots were generated. In these plots, the x-axis represented the nuclear volume, while the y-axis indicated the overlapped volume ratio of surface_Nuclei to surface_YAP.

To calculate the ratio of microchannels in the total 3D hydrogel volume, Z-stack scans of fluorescent-labeled microfilaments were reconstructed using the same method. The ratio of microchannels (*P*) was determined by the formula: 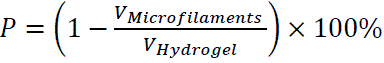.

### RNA extraction and quantitative reverse transcription polymerase chain reaction (qRT-PCR)

The RNA extraction of hydrogel constructs was performed following the previous work. Briefly, the cell-laden hydrogel samples were transferred into 500 µL of NucleoZOL (Macherey-nagel, 1506/001) after 2 weeks of incubation. The tissue samples were homogenized and then incubated at 25 °C for 10 min. DNase-free water was added to the mixture, then centrifugated for 10 min at 12000 rcf. The supernatants were mixed with membrane binding solution and isopropanol and then purified using ReliaPrep RNA Clean-up kit (Promega, Z1073). The yield and purity of the isolated RNA were determined with a spectrophotometer (Thermo Scientific, NanoDrop One). An A260/280 ratio of between 1.8-2.1 was accepted as adequate quality for the RNA samples. The isolated RNA was transcribed to complementary DNA following the instruction of GoScript Reverse Transcriptase kit (Promega, A5003). The relative gene expression levels were determined on real-time PCR System (Applied Biosystems, QuantStudio 3) with the GoTaq qPCR kit (Promega, A6001). The GAPDH housekeeper gene was used as an internal control for the normalization of RNA levels. Primer pairs used in this study are available in the supporting information (**Table S4**).

### Statistical analysis

Statistical analysis was carried out using GraphPad Prism (x64, v. 9.6.0). Data was analyzed using unpaired *t*-test and presented as mean ± SD unless stated otherwise. Alpha was set to 0.05 and differences between two experimental groups were judged to have statistical significance at *p < 0.05, **p < 0.01, ***p < 0.001, and ****p<0.0001; ns represents “no significant difference” between two groups.

## Acknowledgements

The authors express their appreciation to the donors who generously provided their tendon tissues for this study. The authors gratefully acknowledge clinical teams at Balgrist University Hospital and the Laboratory for Orthopedic Biomechanics for providing human tendon tissue samples/tenocytes. This work was done within the framework of the ALIVE initiative (Advanced Engineering with Living Materials) and funded by the SFA-AM program (Strategic Focus Area – Advanced Manufacturing). The ScopeM are thanked for their support on confocal imaging. H. L. thanks Jakub Janiak for his kind assistance with the figure illustration. H. L. acknowledges Amelia Hasenauer for her kind help with hydrogel imaging.

## Data Availability Statement

The data that support the findings of this study are available from the corresponding author upon reasonable request.

