## Supplementary files for "Filamented Light (FLight) Biofabrication of Mini-Tendon Models Show Tunable Matrix Confinement and Nuclear Morphology"

### Title

### Content

Figure S1 – S9

Table S1 – S4

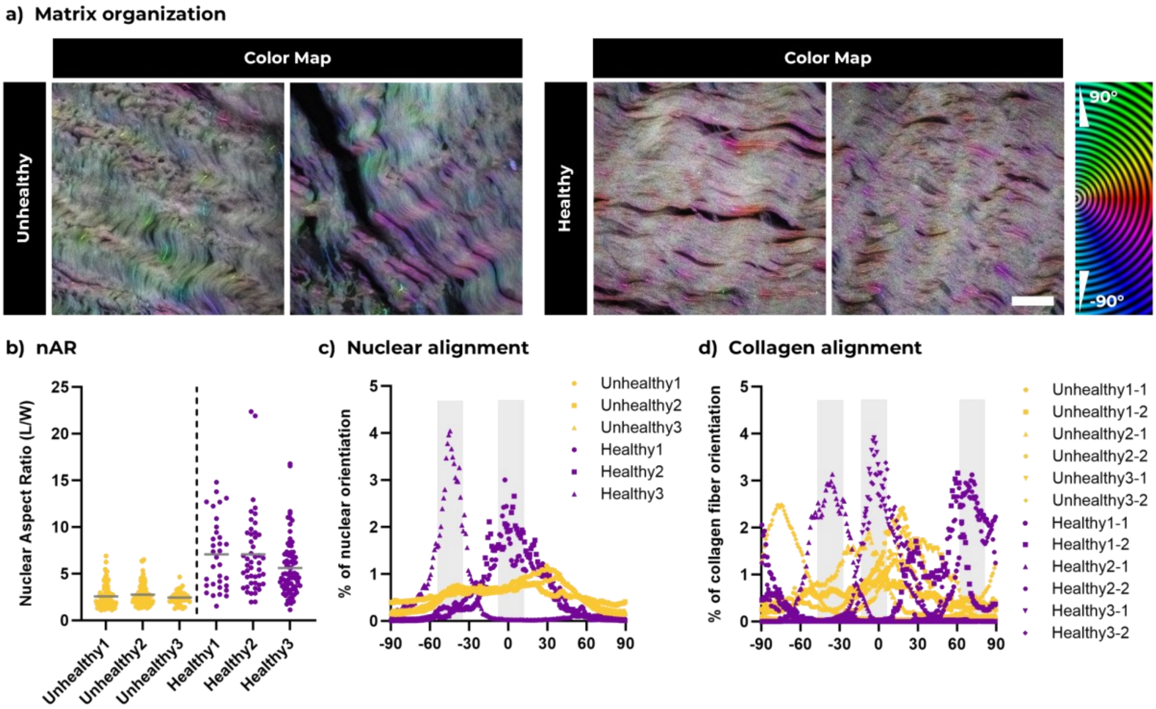


**Figure S1. Nuclei morphology, orientation, and collagen type I fiber alignment in tendons from different patients**. **a**) Orientation color maps generated using the OrientationJ plugin, corresponding to the confocal microscopy images. Scale bars: 20 µm. **b**) Nuclei aspect ratio (nAR) of tenocytes in both healthy and unhealthy tendons and **c**) distribution of nuclear orientation, determined by the direction of the long axis of nuclei. Gray boxes highlight the angle ranges identified as aligned collagen. **d**) Orientation distribution of collagen fibers in healthy versus unhealthy human tendons. Gray boxes highlight the angle ranges identified as aligned collagen.

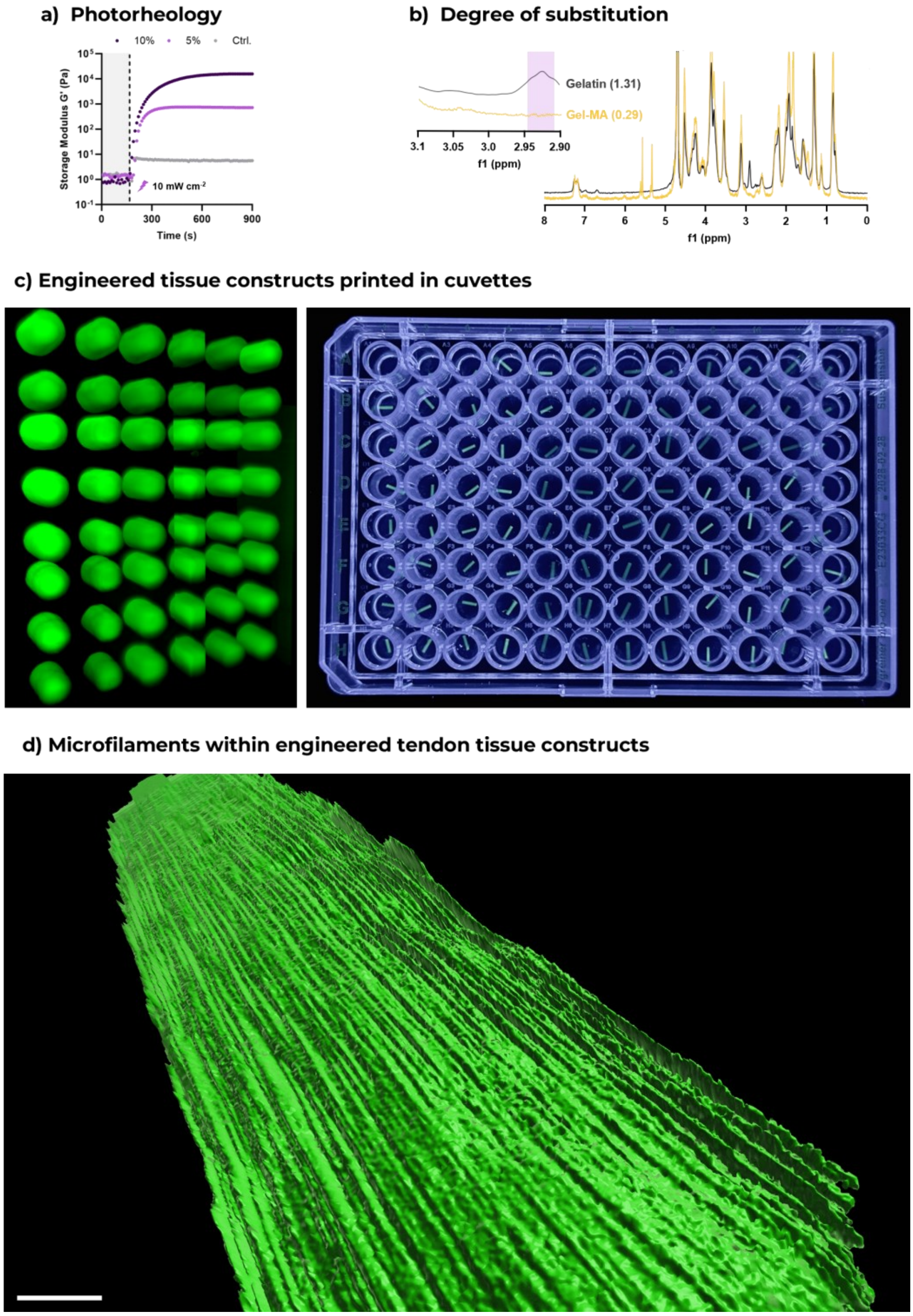


**Figure S2. Characterization of Gel-MA photoresin.** **a**) Photorheological analysis of the photoresin used in this study, conducted under 405 nm irradiation at an average intensity of 10 mW cm$^{-2}$. **b**) Comparative $^{1}$H-NMR spectra of Gelatin and Gel-MA, measured in $D_2O$. **c**) Photographs of FLight hydrogel constructs after being transferred to a 96-well plate. 96 constructs were printed within 12 seconds using fluorescent-labeled Gel-MA. **d**) 3D reconstruction of FLight hydrogel constructs from light-sheet imaging using Imaris software, which highlights the continuous microfilament within the matrix. Scale bar: 200 µm.

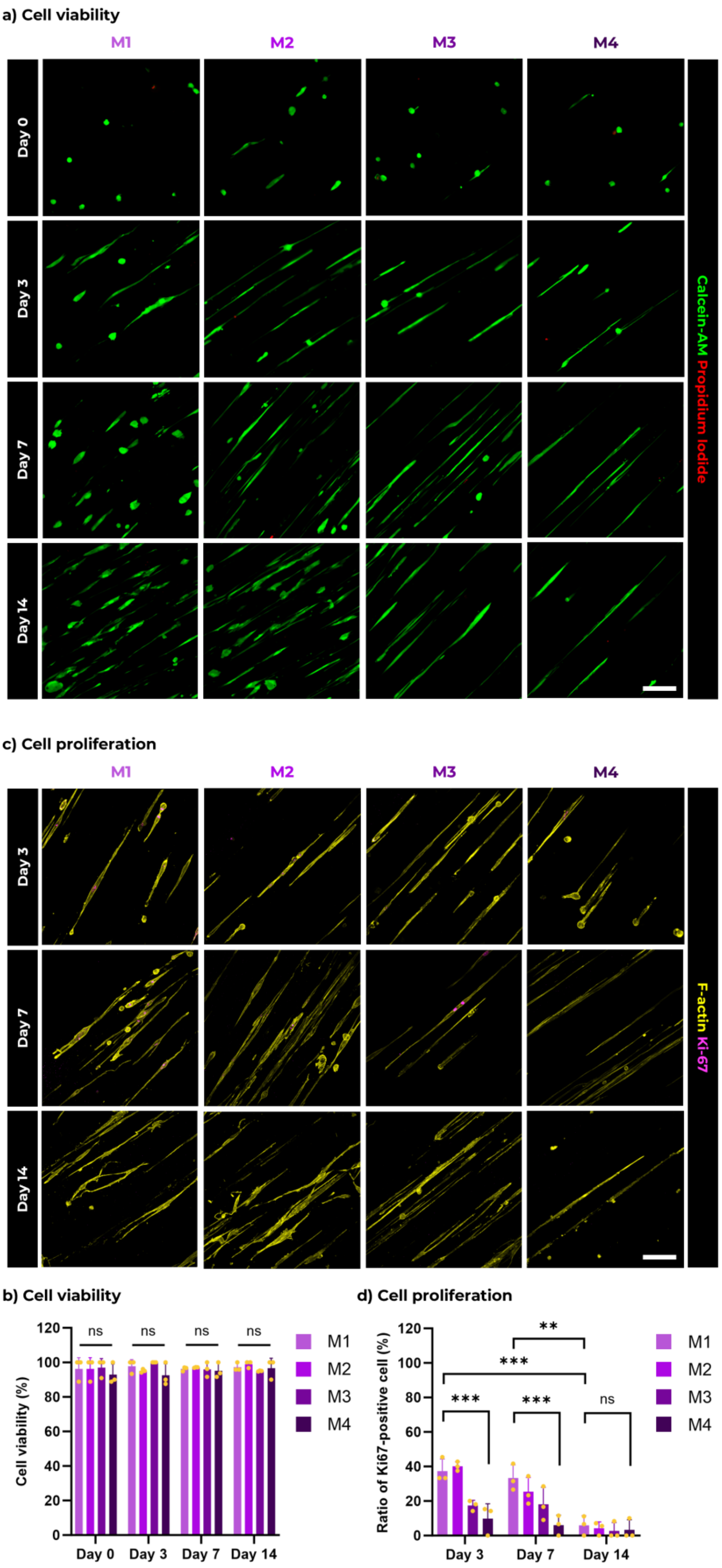

a) Cell viability
M1
M2
M3
M4
Day 0
Day 3
Day 7
Day 14
Calcein-AM Propidium Iodide
c) Cell proliferation
M1
M2
M3
M4
Day 3
Day 7
Day 14
F-actin Ki-67
b) Cell viability
Cell viability (%)
0
20
40
60
80
100
120
ns
ns
ns
ns
Day 0
Day 3
Day 7
Day 14
M1
M2
M3
M4
d) Cell proliferation
Ratio of Ki67-positive cell (%)
0
20
40
60
80
100
120
**
***
***
***
ns
Day 3
Day 7
Day 14
M1
M2
M3
M4

**Figure S3. Cell viability and proliferation in FLight matrices. a**) Panel shows representative confocal Live/Dead images of encapsulated cell in different FLight matrices after 0, 3, 7, and 14 days of culture and **b**) quantitative analysis of cell viability. **c**) Representative immunofluorescent confocal images of Ki-67 expression in human tenocytes and **d**) quantitative analysis of Ki-67 positive cell ratio, depicting the cell proliferation in different FLight matrices. Scale bar: 20 μm.

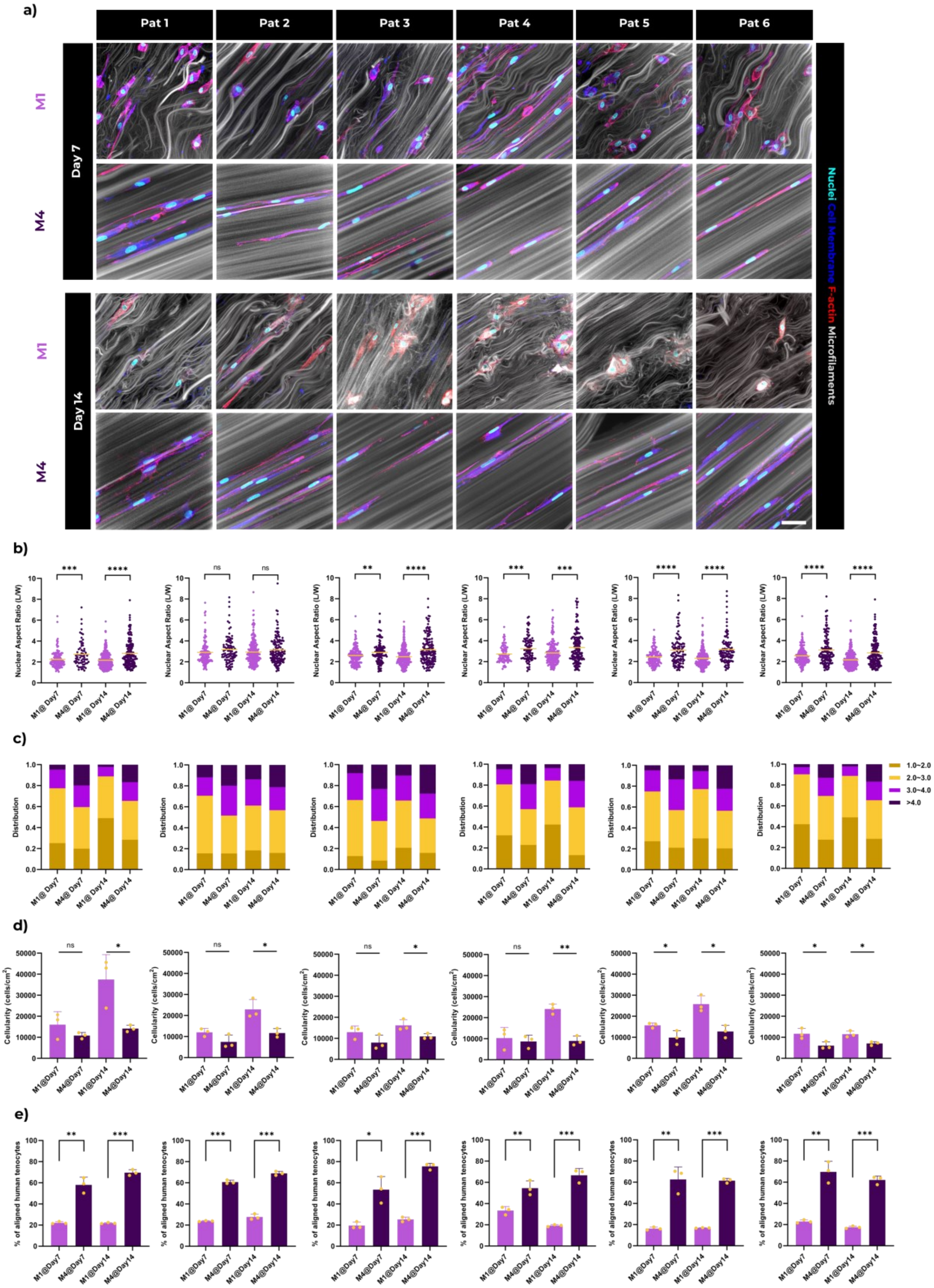


**Figure S4. Nuclei morphologies in mini-tendon constructs using patient-specific tenocytes. a)** Representative confocal microscopy images showing patient-specific tenocytes in different mini-tendon constructs after 7 and 14 days of culture. Scale bar: 50 µm. **b)** analysis of nuclei aspect ratio (nAR) in patient-specific tenocytes and **c)** distribution of nAR in matrix1 and 4. **d)** Cellularity comparison between Matrix 1 and Matrix 4 at day 7 and day 14. **e)** Percentage of aligned human tenocytes in different mini-tendon constructs, measured after 7 days (left) and 14 days (right) of incubation. The aligned cells are characterized by an angle deviation of f-actin from the projection direction between -10 to 10°.

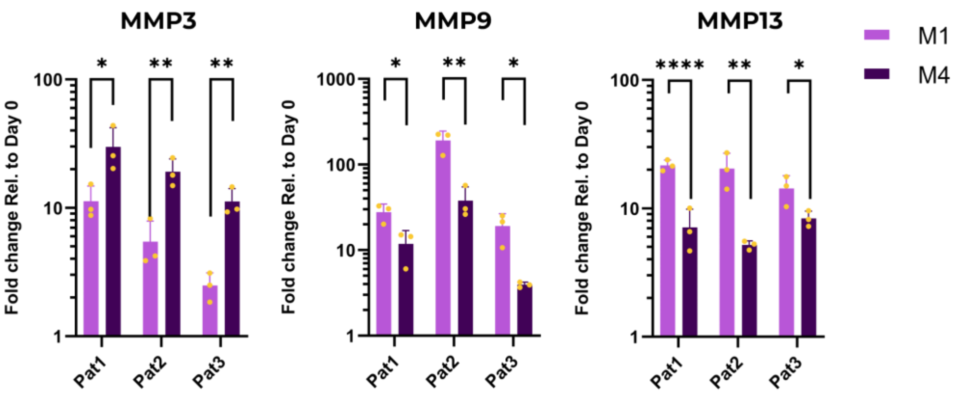


**Figure S5.** Gene expression of encapsulated human tenocytes in various matrices after 14 days of culture, normalized to Day 0. Refer to Table S5 for details on primer designs.

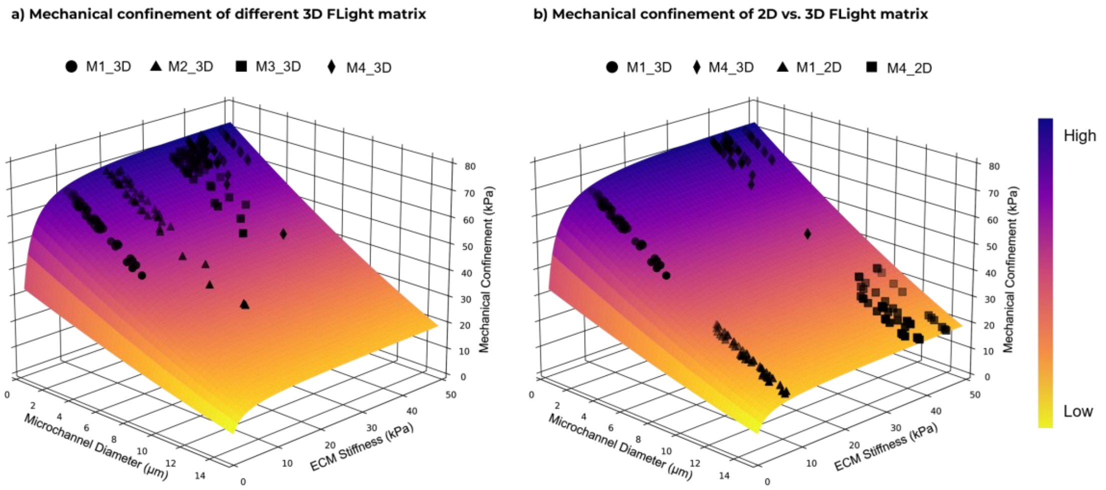


**Figure S6.** A model[1] of mechanical confinements applied by microarchitectures when **a**) cells are present in microchannels of varying filamented matrices vs. **b**) cells adhered on the surface of FLight hydrogel.

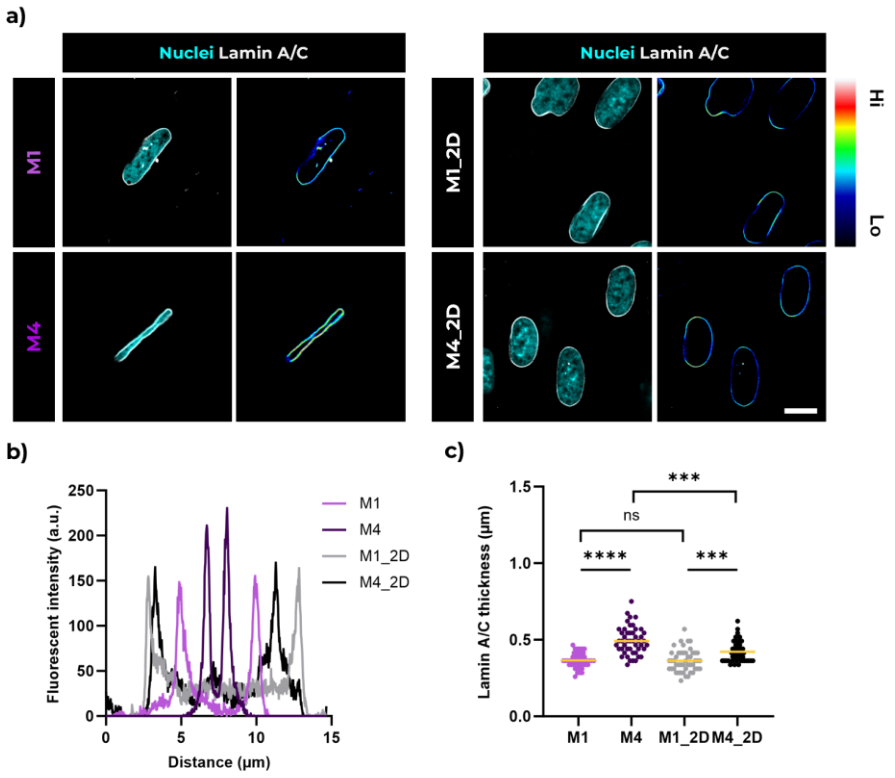


**Figure S7. Nuclear deformation and lamin A/C distribution in human tenocytes. a**) Representative confocal microscopy images indicating the variability in the spatial distribution of lamin A/C within tenocytes in M1, M4 hydrogel matrices, and on the 2D surface of FLight hydrogel constructs. Images are cross-sectional views of nuclei. Scale bar: 10 µm. **b**) Example of lamin A/C distribution cross the short axis of nuclei shown in a), using data from 'Plot Profile' function in Fiji. **c**) Comparative analysis of lamin A/C thickness in tenocytes under different conditions (n=3, dataset size=48).

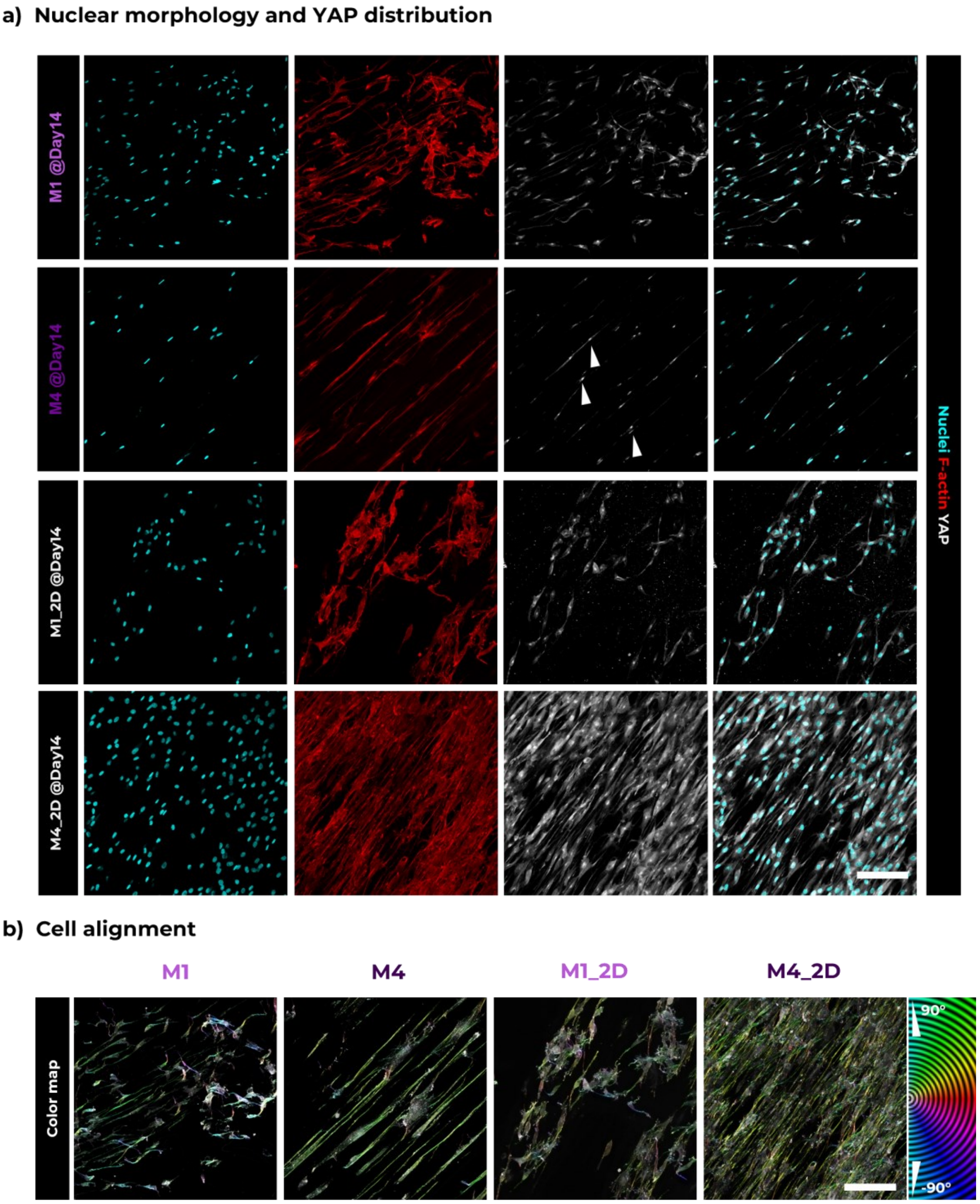


**Figure S8. Nuclei morphology and YAP nuclear activation of tenocytes under different mechanical confinements. a**) Representative confocal microscopy images (20x) of human tenocytes within various mini-tendon constructs (Matrix 1 and 4), and on the 2D surface of FLight hydrogel constructs (Matrix 1 and 4) after 14 days of culture. Scale bar: 200 µm. **b**) Orientation color maps of F-actin generated by the OrientationJ plugin.

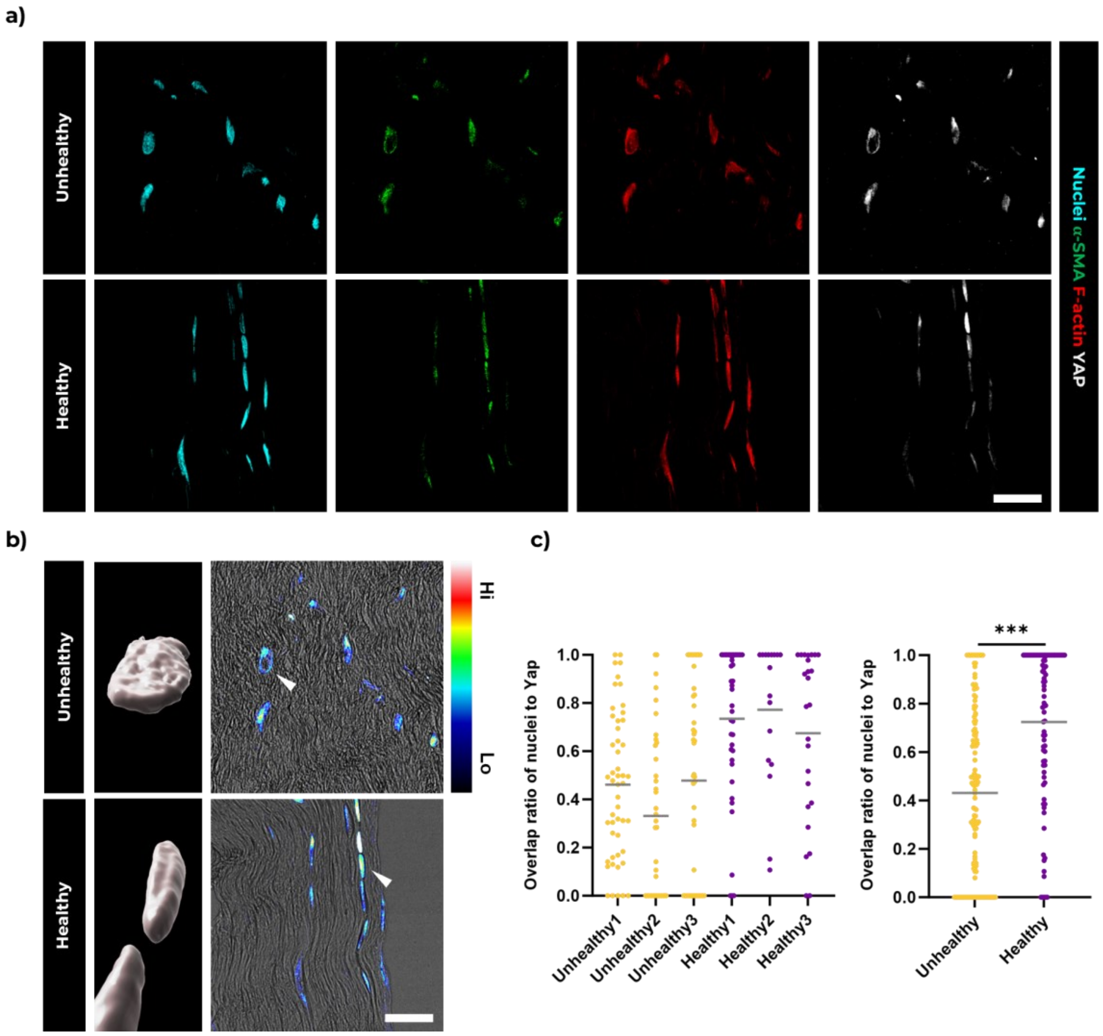


**Figure S9. YAP nuclear activation in healthy and unhealthy human tendons. a**) Representative confocal microscopy images showcasing human tendons. Scale bar: 20 µm. **b**) 3D reconstruction of nuclei and YAP intensity distribution map. The arrows highlight the nuclei reconstructed using Imaris 9.9.0 software, with relative intensity represented on a rainbow scale. Scale bar: 20 µm. **c**) Comparison of the volume overlap ratio of nuclei to nuclear YAP in healthy and unhealthy human tendons, analyzed from 6 patients with a dataset size > 82.

| **Patient ID (#)** | **Age** | **Tissue** | **Diabetes** no = 0; yes = 1 | **Sex** male = 1; female = 2 |
|---|---|---|---|---|
| 106 | 16 | Semitendinosus (right) | 0 | 1 |
| 152 | 47 | Semitendinosus (right) | 0 | 1 |
| 154 | 17 | Semitendinosus (right) | 0 | 2 |
| 763 | 27 | Semitendinosus (right) | 0 | 2 |
| 854 | 51 | Semitendinosus (right) | 0 | 1 |
| 1206 | 31 | Semitendinosus (right) | 0 | 2 |

**Table S1. Comprehensive patient information and details on human tenocytes employed in this study.**

| Parameter | Meaning | Value | Origin |
|---|---|---|---|
| $E_c$ | Cell stiffness | 1000 Pa | [2] |
| $D_c$ | Normal cell diameter (Day0) | 12.5 μm | Measured |
| $V_c$ | Poisson’s ratio of the cell | 0.55 | Assumed |
| $E_{ECM}$ | ECM stiffness | 1-50 kPa | Measured |
| $V_{ECM}$ | Poisson’s ratio of the extracellular matrix | 0.3 | Assumed |
| R | Cell radius | 6.25 μm | Measured |
| $W_{Track}$ | Width of microchannels | 1-10 μm | Measured |

**Table S2. Parameters used in the model depicted Figure S6.**

| Resources | Manufacturer | Catalog | Host Species | Dilution |
|---|---|---|---|---|
| | | **Primary antibodies** | | |
| Anti-Collagen I | abcam | ab138493 | Rabbit | 1:200 |
| Anti-Smooth Muscle Actin | Thermo Fisher Scientific | 14-7960-82 | Mouse | 1:200 |
| Anti-YAP1 | proteintech | 13584-1-AP | Rabbit | 1:100 |
| Anti-Lamin A/C | Thermo Fisher Scientific | MA5-35284 | Rabbit | 1:200 |
| Anti-Ki-67 | BD Biosciences | 550609 | Mouse | 1:200 |
| | | **Secondary antibodies & Dyes** | | |
| Hoechst 33342 | Thermo Fisher Scientific | 62249 | - | 1:1000 |
| Phalloidin-Tetramethylrhodamin B | Sigma | P1951 | - | 1:1000 |
| CellMask Deep Red | Thermo Fisher Scientific | C10046 | - | 1:1000 |
| Goat anti-Mouse IgG(H+L), Alexa 488 | Invitrogen | A-11001 | - | 1:500 |
| Goat anti-Rabbit IgG(H+L), Alexa 647 | Invitrogen | A-21244 | - | 1:500 |

**Table S3. Antibodies/dyes for (Immuno)fluorescence staining and confocal microscopy imaging.**

| Gene | Name | Forward Primer | Reverse Primer |
|---|---|---|---|
| MMP3 | Matrix metalloproteinase 3 | ATTCCATGGAGCCAGGCTTTC | CATTTGGGTCAAACTCCAACTGTG |
| MMP9 | Matrix metalloproteinase 9 | GATGCGTGGAGAGTCGAAAT | CTATCCAGCTCACCGGTCTC |
| MMP13 | Matrix metalloproteinase 13 | CATGAGTTCGGCCACTCCTT | AAGTGGCTTTTGCCGGTGTA |

**Table S4. Primers for qRT-PCR.**
